# Cellular heterogeneity in the response of *Burkholderia thailandensis* to antibiotics and phages in synthetic spatial refuges

**DOI:** 10.64898/2026.09.26.754594

**Authors:** Samuel Kraus, Megan L Fletcher, Urszula Łapińska, Sunee Korbsrisate, Sarah V Harding, Krasimira Tsaneva-Atanasova, Mark A T Blaskovich, Stefano Pagliara

## Abstract

Understanding the complex interactions between phage, antibiotics and bacteria is vital for the development of rationale phage-antibiotic therapies. These interactions have traditionally been investigated via bulk assays, thus masking eventual cellular heterogeneities and the impact of spatial refuges on phage and antibiotic efficacy. Using microfluidics-based time-lapse microscopy, we reveal large phenotypic heterogeneity in the response of *Burkholderia thailandensis* to both the bacteriophage ΦBp-AMP1 and sub-inhibitory concentrations of two selected antibiotics ciprofloxacin and trimethoprim. We discover that simultaneous exposure to ciprofloxacin and phage decreases the size of individual *B. thailandensis* cells, whereas phage and trimethoprim together slow down cell doubling. Exposure to ciprofloxacin also causes extended phage lysis of *B. thailandensis* which is preceded by a distinguishable pause in cell elongation, whereas exposure to trimethoprim constrains lysis. As a consequence, exposure to ciprofloxacin further hinders the growth of *B. thailandensis* in the presence of phage, whereas exposure to trimethoprim enhances growth in the presence of phage. These findings help understand the effects of the interactions between antibiotics and phage on bacterial subpopulations in spatial refuges that are notoriously difficult to treat with antibiotics.

## Introduction

*Burkholderia (B.) pseudomallei* is the causative agent of the disease melioidosis that is endemic in at least 79 subtropical and tropical countries (1). *B. pseudomallei* is intrinsically resistant to penicillin, ampicillin, gentamicin, all first- and second generation cephalosporins and polymyxins (2). *B. pseudomallei* can survive and multiply in neutrophils, monocytes and macrophages and commonly forms biofilms, adding to its defensive arsenal (1,3). The current treatment for melioidosis is lengthy, has poor patient compliance with the risk of relapse and no vaccine currently exists (4,5). The concurrent misuse and overuse of antibiotics across medicine and agriculture, as well as rising global temperatures and an increased dissemination of endemic bacteria further aggravate the spread of AMR (6). Concomitantly, climate change and increasingly severe weather events are spreading the endemic boundaries of melioidosis, increasing its clinical burden (7). Recent progress in the development of vaccines, monoclonal antibodies and drug screening for antivirulence therapies have shown promise as novel approaches to combat melioidosis (8–10). However, the risk of adverse drug reactions, low patient adherence to prolonged oral therapy and limited alternatives to the gold-standard antibiotic therapy remain significant challenges (5). Additionally, although promising, all of these approaches are dose-dependent and unable to evolve to overcome emerging resistance during treatment.

Bacteriophages (or phages) are naturally occurring viruses of bacteria that are auto-dosing and evolve to overcome emergent bacterial defences in real-time (11,12). As the most abundant biological entity on earth, phages represent a vast reservoir of naturally occurring antimicrobials (13). Driven by a billion-year arms race between phage and their hosts, bacterial phage resistance against individual phages emerges rapidly (12,14). Phage therapy of melioidosis has shown some promise in improving survival rates in murine models (15) and temperate *B. pseudomallei* phages were recently identified as a potential new avenue for engineered phage-therapy (16), although the rapid emergence of resistance remains a major challenge.

One approach to prevent the emergence of bacterial phage resistance is phage-antibiotic combination treatment that can overwhelm bacterial defences (17). Indeed, the combination of phage and antibiotics can create a lose-lose scenario for the bacterial host, achieving a higher bacterial killing than both respective monotherapies (17,18), including in clinical settings (19,20). Although promising, phage-antibiotic interactions are difficult to predict and antibiotics that share the same cellular target may interact differently with the same phage (21).

The current orthodoxy to study the efficacy of phage and phage-antibiotic combinations against bacteria focuses primarily on bulk-culture. While this is certainly relevant, it does not monitor subpopulational heterogeneities such as those encountered in spatial refuges, physical safe zones where bacteria are less exposed to treatment, that are common in nature, for instance in biofilms (22). Microfluidics-based time-lapse microscopy enables the observation of bacterial responses to stressors at the single-cell level in synthetic refuges in real-time (23). Recently, we used our microfluidic ‘mother machine’ platform to show that *Escherichia coli* cells double while shrinking in response to high ciprofloxacin stress (24), that subpopulational phenotypic variants of *E. coli* and *Pseudomonas aeruginosa* survive interaction with antimicrobial peptides through enhanced efflux (25) and that subpopulational heterogeneity emerges in CRISPR-immune *P. aeruginosa* in response to phage DMS3*vir* predation (26).

Using bulk assays, we have also recently shown that the bacteriophage ΦBp-AMP1 (27–29) displays additivism with ciprofloxacin and other quinolones, as well as tetracyclines and beta-lactams when treating *B. thailandensis* in liquid culture (14). Here, we utilise our microfluidics-based time-lapse microscopy platform to dissect the subpopulational response of *B. thailandensis* to phage ΦBp-AMP1, and phage-antibiotic combinations in synthetic refuges mimicking those encountered in naturally structured environments.

## Materials and Methods

### Bacterial culturing

*Burkholderia thailandensis* strain E264 (NC_007650/51 (30)), was stored at - 80 C and streaked onto Lysogeny broth (LB, 10 g/L Tryptone, 5 g/L Yeast extract, 10 g/L NaCl, Melford) agar plates (10 g/L, 1.5% Agar) every two weeks. Overnight cultures were setup by inoculating a single *B. thailandensis* colony from a plate into flasks containing 50 mL of LB broth and were grown for 17 h at 37 °C on shaking platforms set at 200 rpm.

### Propagation and titration of phage

Phage ΦBp-AMP1 (29) was propagated as previously reported (31). Briefly, overnight cultures of *B. thailandensis* E264 were diluted 1000× in 50 mL Luria-Bertani (LB) broth to obtain a bacterial concentration of approximately 2×10^6^ CFU mL^-1^. These sub-cultures were then incubated for 4 h at 37 °C and 200 rpm, allowing them to reach early exponential phase (∼8×10^6^ CFU mL ^-1^). The sub-cultures were infected with ΦBp-AMP1 at an MOI = 0.01 and incubated for 17 h at 37 °C and 200 rpm. The following day, cells were pelleted by centrifugation for 40 minutes at 3000 *g* and the phage-containing supernatant was filtered twice (Sartorius Minisart^TM^ 0.2 µm, Cat# 10730792) to obtain a phage stock. The phage concentration within this stock was then determined via the double agar overlay technique (32). Briefly, LB top agar (10 g/L, 0.5% Agar) was melted in a microwave and allowed to cool to below 40 °C. An aliquot of an overnight culture of *B. thailandensis* was added to the melted top agar at a volume: volume concentration of 1:50. LB agar plates were subsequently coated in a thin layer of the top agar creating a continuous bacterial lawn. In parallel, a 10-fold dilution series of the phage stock was prepared in LB broth. The phage was then titred using the standard double agar overlay technique (33). All plates were incubated at 37 °C for 17 h after which the phage induced plaques in the bacterial lawn were counted and the plaque forming units (PFU) per mL^-1^ of the phage stock was calculated. Phage stocks were stored at 4 °C for a maximum of two weeks before a new propagation was performed. Prior to each use, the phage concentration within the phage stock was determined via the double agar overlay technique described above.

### Single-cell microfluidics

To measure the growth of individual bacteria in real-time we used the microfluidic mother machine device as previously described (34). Briefly, the device is comprised of polydimethylsiloxane replica obtained by molding against a silicon-photoresist master (35) and is made of a central chamber that measures 25 µm and 100 µm in height and width, respectively, and six thousand lateral side channels, each 1 µm in width and height and 25 µm in length (36). Bacteria were prepared by pelleting an overnight culture of *B. thailandensis* for 15 minutes at 3000 *g*. The resulting bacterial pellet was resuspended at a previously optimised OD_600_ of 50 (37), in media obtained by double filtering the supernatant from the centrifuged culture using 0.22 µm filters (Sartorius Minisart^TM^ 0.2 µm, Cat# 10730792). These bacteria were then introduced into the central chamber of the mother machine device from where they reached the lateral channel at an average concentration of one bacterium per channel (38). Next, the device was mounted onto an inverted microscope (IX73 Olympus, Tokyo, Japan) located in a temperature-controlled chamber at 37 °C. Fluorinated ethylene propylene tubing (1/32" × 0.008"), was connected to the device as inlet and outlet tubes connected to a computerised pressure-based flow control system (MFCS-4C, Fluigent) (39). Next, LB broth or LB broth containing 2×10^8^ PFU mL^-1^ phage, 2×10^8^ PFU mL^-1^ phage and ciprofloxacin (Sigma-Aldrich, Cat# 17850) or trimethoprim (Sigma-Aldrich, Cat# T7883) at 0.25× their bulk MIC (14), was continuously supplied into the device at a constant flow rate of 100 µl/h for 9 h, unless otherwise stated.

Simultaneously, bright field images of 20 areas of the mother machine, each containing 23 lateral channels were acquired at 2 min intervals via a 60× 1.2 N.A. objective (UPLSAPO60XW, Olympus) and an sCMOS camera with an exposure time of 0.01 s (Zyla 4.2, Andor, Belfast, United Kingdom) controlled via Labview (40). At the end of each experiment, free nucleotides were stained with Propidium Iodide (PI) Briefly, 20 µM of PI in LB was flowed through the chip at 100 µl/h for 15 minutes, prior to imaging with a Tetramethylrhodamine isothiocyanate (TRITC) filter, a green LED at 100% intensity and a camera exposure time of 0.01 s.

### Image and data analysis

All images were analysed using the ImageJ software as previously described (41,42). Dividing cells were defined as dividing more than three times over the nine-hour incubation period, slowly-dividing cells as dividing twice over the nine-hour incubation period, non-dividing cells as not dividing or dividing once. Filamentation was defined as a cell measuring more than twice the average length of untreated bacteria (i.e. more than 8 µm). Finally, lysis was defined as a visible, physical bursting of cells during 2-minute interval imaging and was confirmed by propidium iodide staining at the end of each experiment. For each experimental condition tested, data were 100 synthetic refuges collated from triplicate experiments in the mother machine, unless otherwise specified. Statistical significance was assessed via unpaired t-tests with Welch’s correction against control experiments, i.e. incubation in LB broth or treatment with phage only, as specified in each figure legend. Data were plotted and analysed using Prism GraphPad v8.3.

## Results

### *B. thailandensis* growth in synthetic refuges

We set out to measure the regrowth of *B. thailandensis* from stationary phase when incubated in LB broth supplied to the lateral channels of the microfluidic ‘mother machine’ (38), which mimic refuges encountered in naturally structured environments. We found that *B. thailandensis* cells settle and then elongate and divide in these synthetic refuges during incubation in LB medium (Figure 1A-C). Specifically, individual *B. thailandensis* cells began elongating after an average of approximately 30 minutes in the structured environment of the ‘mother machine’ device and mid-log phase was reached after 240 min of incubation with an average of 4 cells per channel (filled squares in **Error! Reference source not found.**D). This is in accordance with incubation in well-mixed flasks containing 50 mL of LB medium followed by the colony forming unit (CFU) assay, where mid-log phase was reached after 240 min (empty squares in Figure 1D). The maximum capacity of 7 cells per channel was reached approximately 360 min into the experiment with further progenies pushed out into the main delivery chamber.

**Figure 1.**
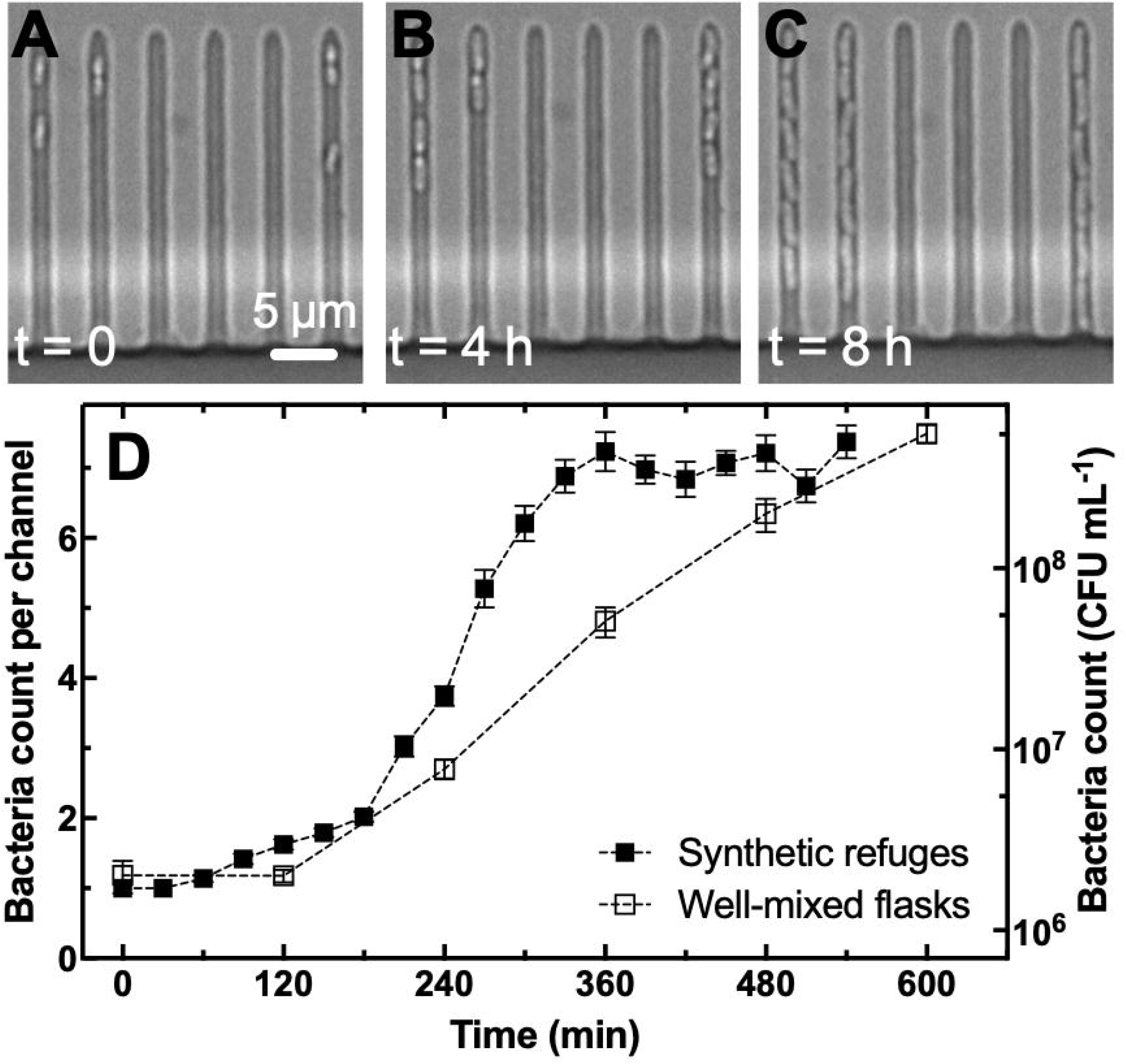
*B. thailandensis* growth in synthetic refuges and in well-mixed flasks. (A-C) Representative images of individual *B. thailandensis* bacteria growing in the lateral channels of the microfluidic ‘mother machine’ after 0h, 4h and 8h of incubation in LB medium supplied at 100 µl h^-1^ and at 37°C starting from a stationary phase inoculum. (D) Corresponding temporal progression of the average number of cells per channel (filled squares) compared to temporal progression of colony forming units per ml counted from *B. thailandensis* growing in well-mixed flasks (empty squares). Symbols and error bars are means and standard errors of the means of cell counts within 100 channels from triplicate experiments in the mother machine or means and standard errors of CFU assays performed in biological triplicate. Some error bars cannot be visualised due to overlap with the datapoints. Dashed lines are guides for the eye.

### Real-time monitoring of antibiotic driven growth inhibition of *B. thailandensis* in synthetic refuges

Next, we inoculated *B. thailandensis* cells taken from stationary phase in synthetic refuges supplied for 540 min with LB medium containing ciprofloxacin at either 0.25× (i.e. 0.5 µg ml^-1^). or 1× MIC (i.e. 2 µg ml^-1^) against *B. thailandensis.* We chose ciprofloxacin as a model antibiotic, as we previously found that this antibiotic has an additive interaction with the phage ΦBp-AMP1 (14).

We found no statistically significant difference in the number of *B. thailandensis* cells per channel in the untreated *B. thailandensis* culture and the *B. thailandensis* treated with ciprofloxacin at 0.25× its MIC value at the end of the 540 min treatment (black squares and blue circles Figure 2A); or between the untreated *B. thailandensis* and *B. thailandensis* treated with ciprofloxacin at 0.25× its MIC value when grown in well mixed flasks (black squares and blue circles in Figure 2B). However, microfluidics-enabled single-cell analysis revealed that exposure to ciprofloxacin at 0.25× its MIC value prolonged the lag phase, with cells performing their first doubling on average 60 min later.

**Figure 2.**
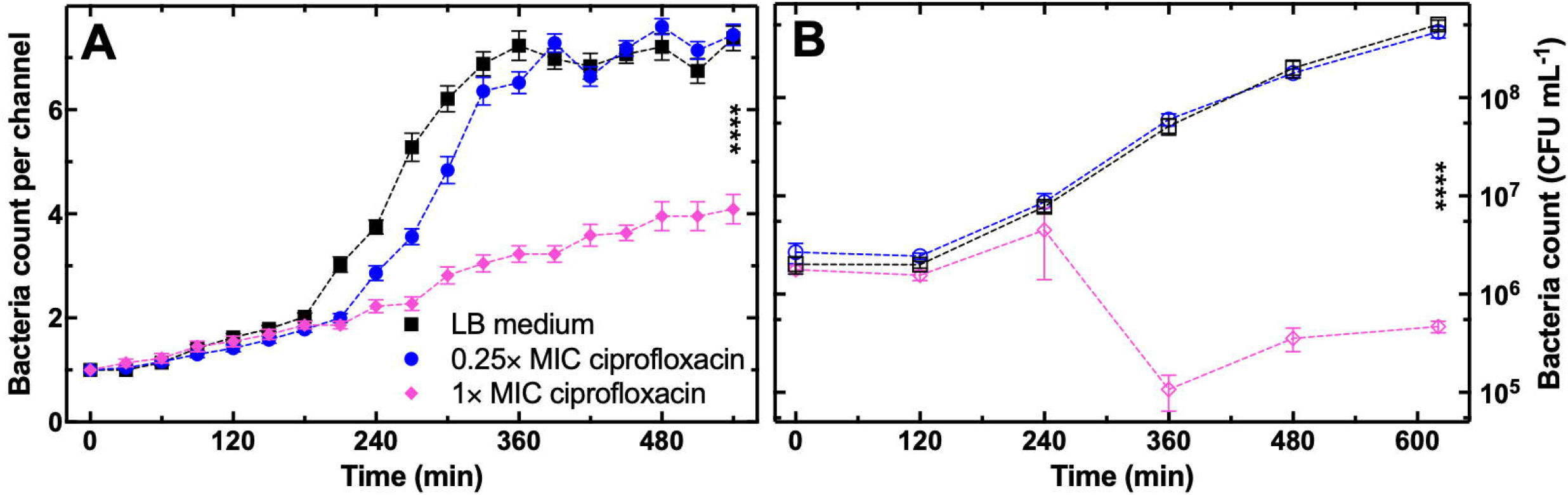
Real-time monitoring of ciprofloxacin driven growth inhibition of *B. thailandensis*. Temporal progression of the average number of *B. thailandensis* bacteria incubated in LB medium (squares), or LB medium with ciprofloxacin at 0.25× (circles) or 1× MIC (diamonds) in (A) synthetic refuges or (B) well-mixed flasks. Symbols and error bars are means and standard errors of the means of cell counts within 100 channels from triplicate experiments in the mother machine or means and standard errors of CFU assays performed in biological triplicate. Some error bars cannot be visualised due to overlap with the datapoints. Dashed lines are guides for the eye. **** indicate a p-value < 0.0001. The data for incubation in LB medium are reproduced from Figure 1D for comparison purposes only.

Increasing the ciprofloxacin concentration to 1× MIC value had a profound impact on *B. thailandensis* growth. Interestingly, we observed different population dynamics in the experiments carried out in the synthetic refuges compared to well-mixed flasks. In synthetic refuges, *B. thailandensis* grew slowly and linearly reaching a final number of cells per channel that was significantly lower compared to untreated *B. thailandensis* (black squares and magenta diamonds in Figure 2A, p-value < 0.0001**Error! Reference source not found.**). In well-mixed flasks, *B. thailandensis* counts did not change during the first 240 min of treatment and then decreased 10-fold, reaching a final number of cells that was significantly lower compared to untreated *B. thailandensis* (black squares and magenta diamonds in Figure 2B, p-value < 0.0001). It is conceivable that the decrease in population in the bulk assay is due to post-antibiotic effect during regrowth on agar plates, where ciprofloxacin bound to DNA gyrase causes extensive cell lysis only after ciprofloxacin removal from the environment (i.e. on LB agar plates without ciprofloxacin), when conditions return to being favourable for growth. These observations are in line with the post-antibiotic effect recently reported for enrofloxacin against *Salmonella enterica* (43) and further highlight the importance of real-time single-cell assays, as the one we introduce for *B. thailandensis* in this study, that does not require post-treatment counting on agar plates.

### Phenotypic heterogeneity in *B. thailandensis* during exposure to ciprofloxacin

Next, we set out to investigate phenotypic heterogeneity in *B. thailandensis* during exposure to ciprofloxacin, a phenomenon that cannot be investigated via standard CFU assays but necessitates the development of single-cell assays.

We observed four main *B. thailandensis* phenotypes: ‘fast dividing cells’ that divided more than three times over 540 min experiments; ‘slowly-dividing cells’ that divided twice; ‘non-dividing cells’ that did not divide or divided only once and filamenting cells that measured more than twice the length of untreated bacteria (Figure 3A-D). We did not observe cell lysis during treatment with ciprofloxacin, in accordance with a recent report on the effect of enrofloxacin against *Salmonella enterica* (44).

**Figure 3.**
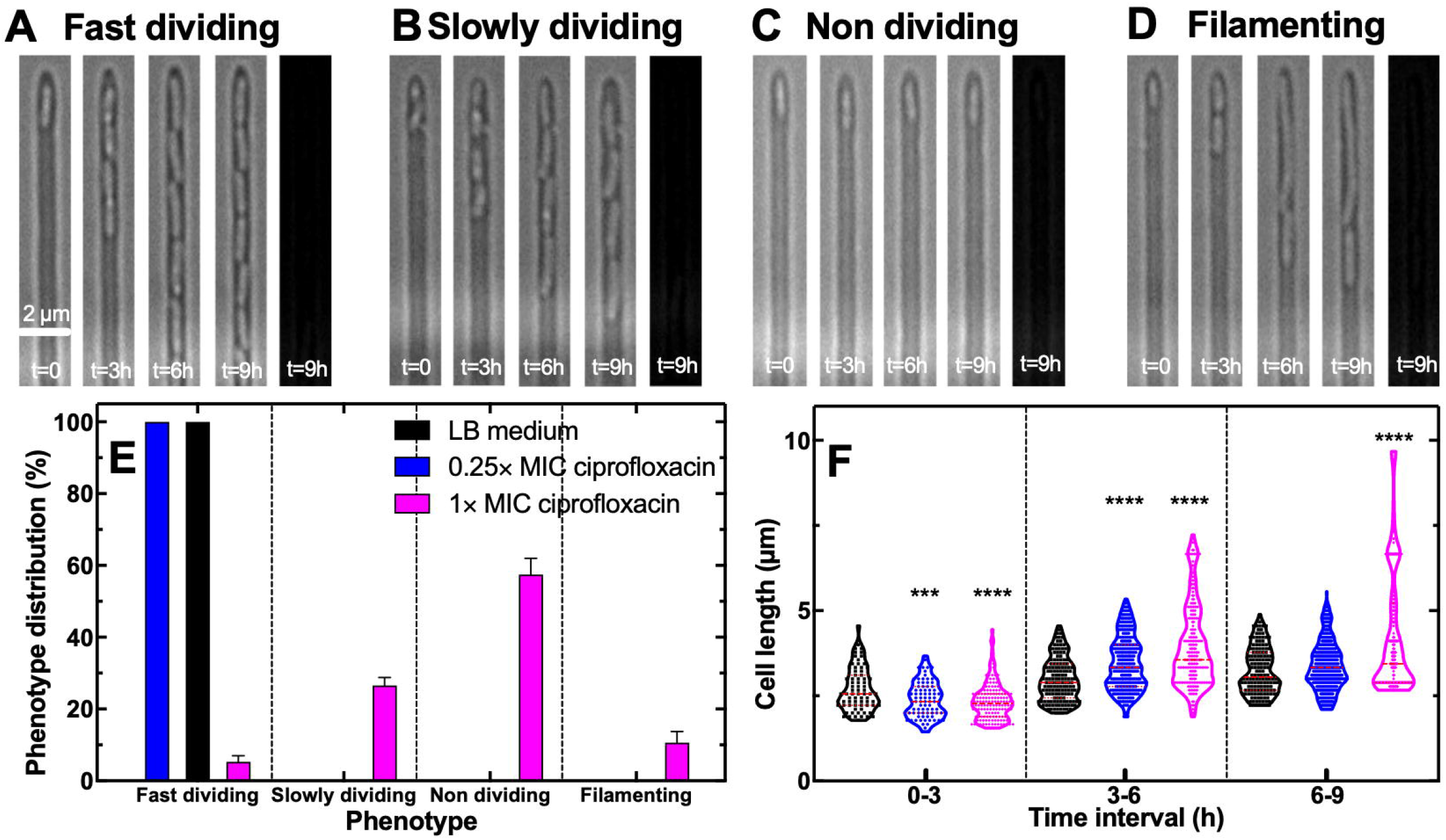
Phenotypic heterogeneity in *B. thailandensis* growth during exposure to ciprofloxacin. (A-D) Representative time-lapse microscopy images of (A) fast dividing, (B) slowly dividing, (C) non-dividing, and (D) filamenting *Burkholderia thailandensis* cells during exposure to ciprofloxacin at 1× MIC. The last image in each panel was acquired in fluorescence mode after supplying the bacteria with propidium iodide. (E) Distribution of each phenotype during incubation in LB medium (black bars), or exposure to ciprofloxacin at 0.25× (blue bars) or 1× MIC (magenta bars). The bar and error bars are the means and standard deviation obtained from averaging over the distributions in individual synthetic refuges. (F) Temporal progression of the cell length of individual *B. thailandensis* cells. Images were taken from five different microfluidic channels from three different experiments to achieve an average of 150 individual measurements in the 0-3 h interval and 300 individual measurements in the 3-6 h and 6-9 h interval, in each condition, respectively. Median, upper, and lower quartile are highlighted as red, dashed lines **** indicate a p-value < 0.0001; *** indicate a p-value < 0.001.

We observed only fast dividing cells in the untreated *B. thailandensis* culture and the *B. thailandensis* culture treated with ciprofloxacin at 0.25× MIC (black and blue bars in Figure 3E). In contrast, during exposure to ciprofloxacin at 1× MIC, the fast-dividing phenotype occurred in only 5% of refuges, the slowly dividing phenotype in 27% of refuges, the non-dividing phenotype in 57% of refuges, and the filamenting phenotype in 11% of refuges (magenta bars in Figure 3E). When we introduced propidium iodide staining at the end of the 540 min experiments, we did not observe any staining in any of the phenotypes, suggesting that all phenotypes maintained an intact membrane. It is conceivable that some cells belonging to the non-dividing phenotype were dead but maintained an intact membrane or were in a viable but non-culturable state.

Our single-cell assay also allowed us to capture changes in *B. thailandensis* cell length during treatment with ciprofloxacin. During the first 3 h in the synthetic refuges, *B. thailandensis* cells displayed a significantly shorter length in increasing ciprofloxacin concentrations, i.e. 2.8 µm, 2.4 µm and 2.3 µm during exposure to LB broth, ciprofloxacin at 0.25× or 1× MIC, respectively (black, blue and magenta plots in Figure 3F, p-value <0.001 via unpaired t-tests with Welch’s correction relative to data for LB medium only). In contrast, between 3 h and 6 h, *B. thailandensis* bacteria became significantly longer at increasing ciprofloxacin concentrations, i.e. 3.0 µm, 3.4 µm and 3.9 µm during exposure to LB broth, ciprofloxacin at 0.25× or 1×MIC, respectively. This trend was maintained in the 6-9 h temporal window, with cells growing significantly longer during 1× MIC ciprofloxacin exposure. i.e. 3.0 µm, 3.4 µm and 4.3 µm (black, blue and magenta plots in Figure 3F).

### Exposure to trimethoprim and phage slows down the doubling of individual *B. thailandensis* cells, whereas ciprofloxacin and phage decrease their size

Next, we set out to quantitatively compare the effect of phage ΦBp-AMP1 and the effect of phage ΦBp-AMP1 in combination with sub-inhibitory concentrations of either ciprofloxacin or trimethoprim on the growth of individual *B. thailandensis* cells. We used these two combinations because in our previous work we found that ciprofloxacin and ΦBp-AMP1 have an additive effect, whereas trimethoprim and ΦBp-AMP1 have an antagonistic interaction (14). In all cases we used ΦBp-AMP1 at a concentration of 2×10^8^ PFU ml^-1^ since we have previously shown that when phage are supplied in the main chamber at this concentration, each synthetic refuge is explored by at least one phage within 2 h (14). Considering that each refuge tipycally hosts one or two cells within this time period, these experimental conditions approximate a nominal multiplicity of infection (MOI) of 1.

We found that exposure to either phage or phage and ciprofloxacin at 0.25× its MIC value did not significantly affect the doubling time of individual *B. thailandensis* cells compared to exposure to LB medium only. In comparison, there was a significant increase in doubling time during exposure to phage and trimethoprim at 0.25× MIC (i.e. 0.25× 32 µg ml^-1^, Figure 4A, p-value < 0.0001 for generations 0-2 and < 0.001 for generation 3 according to t-tests with Welch’s correction with respect to LB medium data).

**Figure 4.**
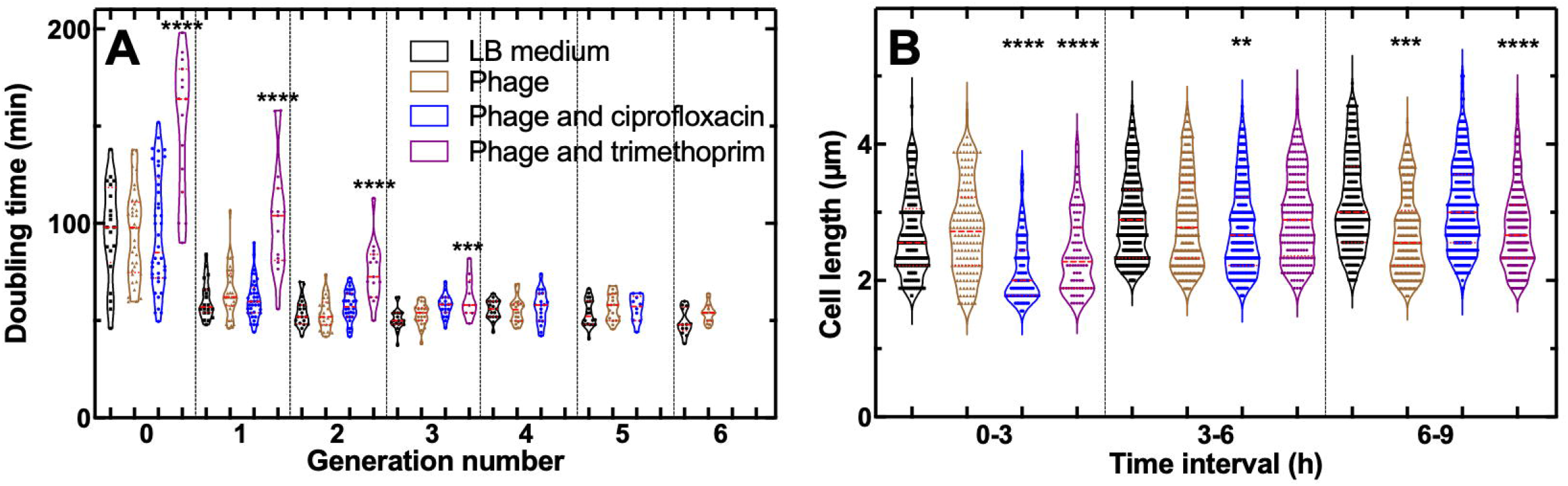
Exposure to ciprofloxacin and phage decrease the size of individual *B. thailandensis* cells, whereas phage and trimethoprim slow down their doubling. (A) Generation time of individual *B. thailandensis* cells during exposure to LB medium (black squares), 2×10^8^ PFU mL^-1^ phage (brown triangles), phage and ciprofloxacin at 0.25× MIC (blue circles), or phage and trimethoprim at 0.25× MIC (purple hexagons). These data were obtained from 15 synthetic refuges from 3 independent experiments. (B) Corresponding temporal dependences of the length of individual *B. thailandensis* cells. Median, upper, and lower quartile are presented as red dashed lines. **** indicate a p-value < 0.0001. *** indicate a p-value of < 0.01.

Moreover, during the first 3 h of exposure to phage and ciprofloxacin, or phage and trimethoprim the length of individual *B. thailandensis* cells was significantly lower compared to exposure to LB medium or phage only (Figure 4B, 0-3, p-value < 0.0001). In the 3-6 h temporal window, only exposure to phage and ciprofloxacin caused a significant decrease in cell size, whereas in the 6-9 h window decrease in cell size was recorded during exposure to phage only and the combination of phage and trimethoprim (Figure 4B), suggesting that the effect of phage and antibiotic treatment on cell size is transient and dynamic.

### Exposure to ciprofloxacin causes extended phage lysis of *B. thailandensis*, whereas exposure to trimethoprim constrains lysis

We also found that some *B. thailandensis* cells lysed in the presence of phage which did not occur in any non-phage containing experiment. Moreover, we did not observe filamentation during exposure to either phage or phage-ciprofloxacin or phage-trimethoprim when these antibiotics were employed at 0.25× their respective MIC values. We determined lysis events by a visual bursting of cells observed during 2-minute interval imaging and confirmed lysis by propidium iodide staining of freed nucleotides at the end of the experiment (Figure 5A, t=9 h). Proximity to the phage source (i.e. the main microfluidic delivery chamber) was not required for initial bacterial lysis as the first lysis event in each refuge could occur in all positions of the synthetic refuges during exposure to phage or phage-antibiotic combinations (Figure 5B).

**Figure 5.**
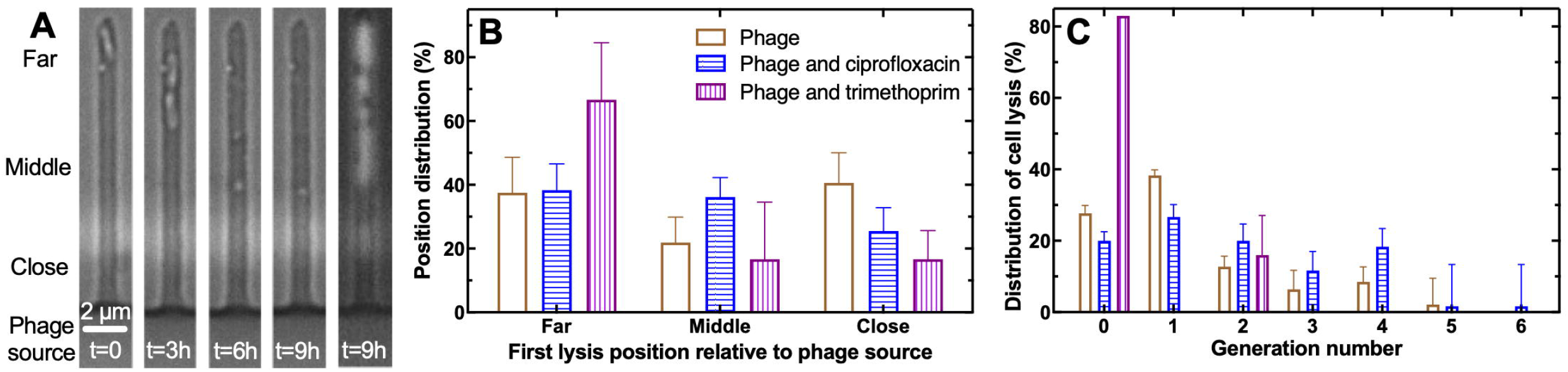
The timing of lysis of individual *B. thailandensis* cells by phage is extended in the presence of ciprofloxacin and reduced in the presence of trimethoprim. (A) Microscopy images illustrating a cell lysis event occurring at the far side of a synthetic refuge with respect to the phage supply source. Far, Middle and Close were defined as one third of the length of the synthetic refuge each, with respect to the phage supply source. (B) Distribution of the position of first cell lysis event within individual synthetic refuges during exposure to 2×10^8^ PFU mL^-1^ phage (brown empty bars), phage and ciprofloxacin at 0.25× MIC (blue horizontally striped bars), or phage and trimethoprim at 0.25× MIC (purple vertically striped bars). (C) Corresponding distribution of the generation number in which cell lysis occurred. Bars and errors were obtained by averaging measurements from 15 synthetic refuges from triplicate experiments.

Ciprofloxacin and trimethoprim had opposite effects on the timing of *B. thailandensis* lysis. Exposure to phage and ciprofloxacin caused sustained lysis across 7 consecutive *B. thailandensis* generations, thus extending the temporal window in which lysis occurred, compared to exposure to phage alone. In contrast, exposure to phage and trimethoprim caused the majority of lysis events in generation 0 and no lysis occurred after generation 2 (Figure 5C).

Next, we set out to investigate whether bacteria that would eventually lyse were measurably different to untreated bacteria. We found that although cells that would eventually lyse displayed cell sizes and doubling times prior to lysis similar to untreated cells (Figure 6A,B), these cells displayed a distinguishable pause in elongation prior to lysis by phage (Figure 6C). Cells that lysed in the first bacterial generation, i.e. during the lag phase of growth, displayed halted elongation for an average of 44 minutes during exposure to phage alone or phage and ciprofloxacin, before eventual lysis (Figure 6C). Halted elongation prior to lysis events in the bacterial generations two, three and four halved to 21 minutes in both treatment conditions (Figure 6C). We did not further investigate lysis during exposure to phage and trimethoprim because the majority of lysis events occurred at generation 0 (Figure 5C).

**Figure 6.**
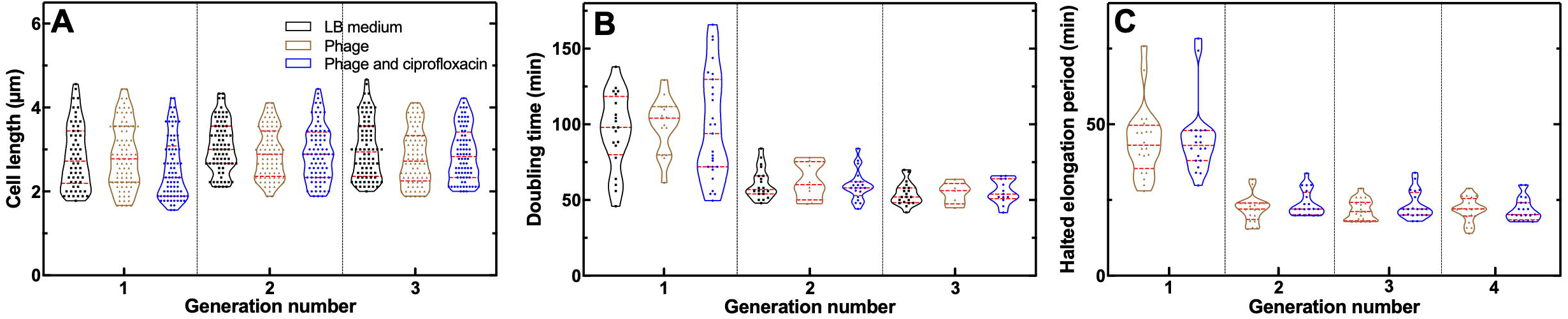
Individual *B. thailandensis* cells stop elongating before lysis. Dependence on generation number for the distribution of (A) cell length and (B) doubling time for *B. thailandensis* cells during exposure to LB medium (black squares) and *B. thailandensis* cells that ultimately lysed during exposure to 2×10^8^ PFU mL^-1^ phage (brown triangles) or phage and ciprofloxacin at 0.25× MIC (blue circles). (C) Corresponding dependence on generation number for the distribution of halted elongation period before lysis. Measurements were obtained from 15 synthetic refuges from triplicate experiments.

### Exposure to ciprofloxacin decreases the frequency of *B. thailandensis* cells that double in the presence of phage, whereas trimethoprim enhances this phenotype

Next, we compared the dynamics in the population size per channel of the dividing and lysing *B. thailandensis* phenotypes during exposure to the different treatments.

During exposure to phage only, the population of dividing cells reached mid-log phase after 180 min and stationary phase after 300 min. Exposure to phage and ciprofloxacin at 0.25× MIC caused a delay in the growth of dividing cells that reached mid-log phase after 300 min and stationary phase after 420 min. This delay was even more striking during exposure to phage and trimethoprim at 0.25× MIC with dividing cells reaching mid-log and stationary phase after 420 min and 540 min, respectively (Figure 7**Error****! Reference source not found.**A). Moreover, cells that eventually lysed grew to a greater extent following phage and ciprofloxacin exposure compared to exposure to phage alone, whereas exposure to phage and trimethoprim had the opposite effect limiting the temporal window in which cells grew before lysing (Figure 7B). Therefore, these data suggest that trimethoprim enhances phage efficacy to a greater extent compared to ciprofloxacin.

**Figure 7.**
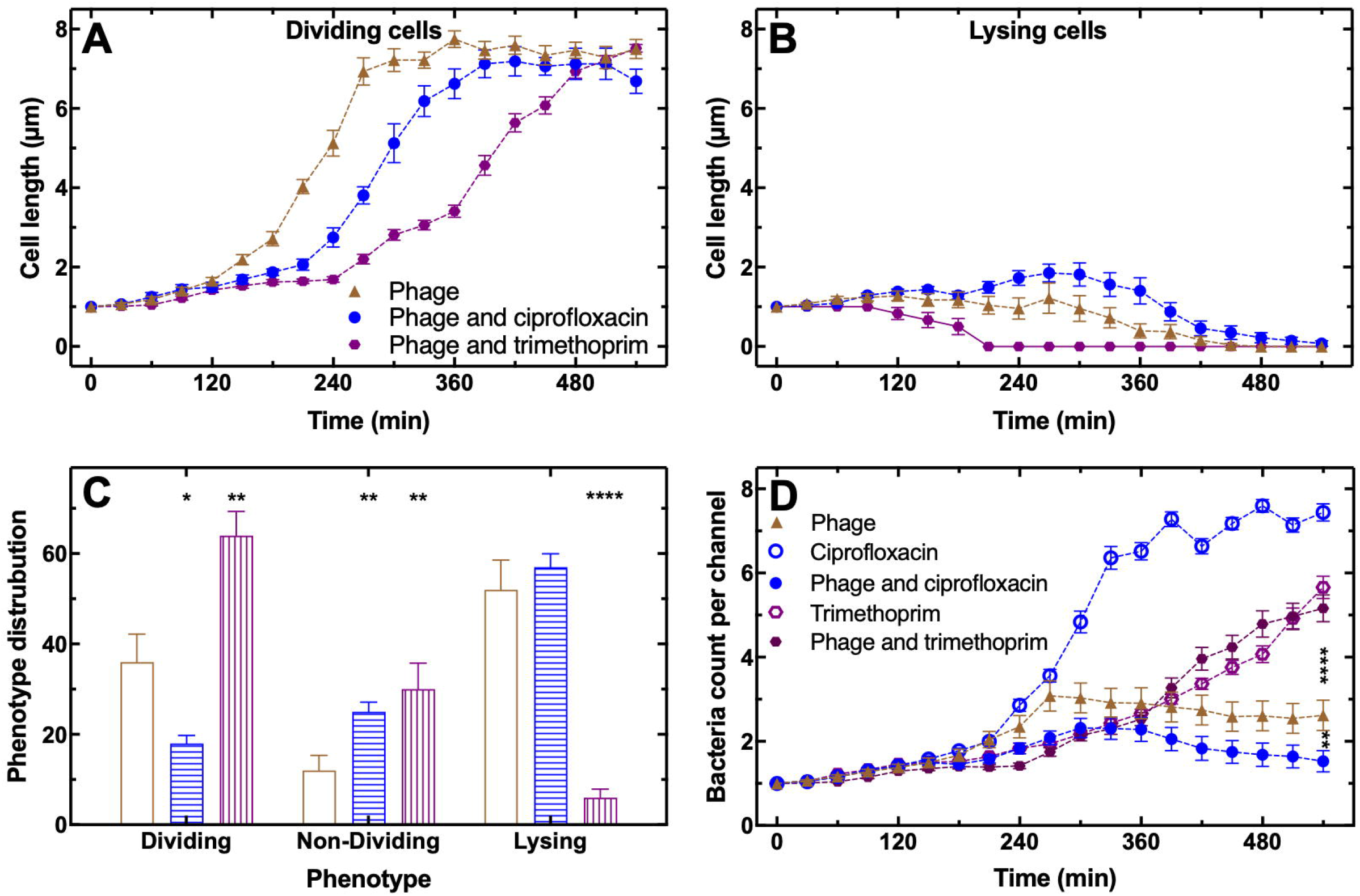
Growth of *B. thailandensis* in the presence of phage is further hindered by ciprofloxacin but enhanced by trimethoprim. Temporal dependence of (A) dividing cells and (B) lysing cells during exposure to 2×10^8^ PFU mL^-1^ phage (brown triangles), or phage and ciprofloxacin at 0.25× MIC (blue circles), or phage and trimethoprim at 0.25× MIC (purple hexagons). (C) Corresponding distributions of the frequency of dividing cells, non-dividing cells and lysing cells. (D) Temporal dependence of the average number of bacteria per channel during exposure to 2×10^8^ PFU mL^-1^ phage (brown triangles), phage and ciprofloxacin at 0.25× MIC (filled circles), phage and trimethoprim at 0.25× MIC (filled hexagons), ciprofloxacin at 0.25× MIC (open circles), or trimethoprim at 0.25× MIC (open hexagons). Points and error bars are the mean and standard error of the mean obtained from measurements in 100 synthetic refuges from triplicate experiments. Some error bars cannot be visualised due to overlap with the datapoints. Dashed lines are guides for the eye. **** indicate a p-value < 0.0001; ** indicate a p-value < 0.01; * indicate a p-value < 0.05.

However, when we compared the distribution of the different *B. thailandensis* phenotypes during exposure to the different treatments, we found that exposure to phage and ciprofloxacin caused a significant decrease in the number of dividing cells, whereas exposure to phage and trimethoprim caused a significant increase in the frequency of this phenotype compared to phage alone (for simplicity, in this analysis we grouped together the fast- and slow-dividing phenotypes introduced above); furthermore, the latter treatment also caused a significant decrease in the frequency of lysing cells. In addition, both antibiotics caused a significant increase in the frequency of non-dividing cells compared to phage alone (Figure 7C, unpaired t-tests with Welch’s corrections performed against the data recorded for incubation in phage alone).

Therefore, when we combined the counts for the different phenotypes over time, we found that exposure to phage and ciprofloxacin significantly decreased the growth of *B. thailandensis* compared to either exposure to phage alone or ciprofloxacin alone. In contrast, exposure to phage and trimethoprim significantly increased the growth of *B. thailandensis* compared to exposure to phage alone, with the dynamics of *B. thailandensis* population size resembling the one recorded during exposure to trimethoprim alone (Figure 7D).

Taken together these data demonstrate the importance of investigating the response of *B. thailandensis* to antibiotics and phage with single-cell resolution. This capability allowed us to understand the previously described phage-antibiotic additivism with ciprofloxacin and antagonism with trimethoprim (14). In the presence of a sub-inhibitory concentration of trimethoprim and phage, a large subpopulation of *B. thailandensis* cells can double albeit at a slow rate and the subpopulation that lyse is reduced compared to phage alone, leading to an antagonistic effect of trimethoprim on phage. In contrast, in the presence of a sub-inhibitory concentration of ciprofloxacin and phage the doubling rate is higher but the subpopulation of cells that doubles is reduced compared to phage alone leading to an additive interaction between ciprofloxacin and phage.

## Discussion

The ability of *B. pseudomallei* to form biofilms is a major risk factor in clinical settings (27) and little is known about phenotypic heterogeneity in the response of this pathogen to antibiotic or phage therapy. Here we introduce an experimental approach that allows to investigate the single-cell response of *B. thailandensis* to antibiotics and phage in synthetic refuges as a tractable model that can simulate the more complex scenario encountered by *B. pseudomallei* cells in spatial refuges in naturally structured environments, such as biofilms (31). Using this approach we show that sub-inhibitory concentrations of the antibiotics ciprofloxacin and trimethoprim extend the lag phase and reduce the length of individual *B. thailandensis* cells, phenotypic responses that can not be captured by traditional microbiology assays. These data are in line with the seminal work of Fridman *et al.* that discovered the ‘tolerance by lag’ strategy in *E. coli* as a response to low antibiotic stress in bulk culture (44).

Furthermore, it has recently been shown that sub-inhibitory concentrations of bacteriostatic antibiotics induce an immediate growth-arrest that is similar to nutrient starvation, whereas bactericidal antibiotics induce a “grow-fast-then-crash” trajectory (45). Our new single-cell data support this hypothesis showing that in the presence of a sub-inhibitory concentration of trimethoprim, bacterial growth is also significantly reduced, whereas ciprofloxacin only prolongs lag phase.

However, ciprofloxacin is also a known driver of cell filamentation that has recently been identified as a key factor for phage-antibiotic treatment of *E. coli* (46). Furthermore, Pons *et al.* recently discovered that filamentous phenotypes of CRISPR-immune *P. aeruginosa* emerged alongside growth arrested phenotypes in response to phage DMS3vir predation (26). In contrast, we found that filamentation did not occur in *B. thailandensis* during exposure to phage ΦBp-AMP1 or a sub-inhibitory concentration of ciprofloxacin, suggesting that this bacterium is more resilient to cell filamentation compared to *E. coli* and *P. aeruginosa* at the concentrations employed in our study.

We also found that *B. thailandensis* was partially resilient to cell lysis by phage ΦBp-AMP1, as it lysed only half of a putatively clonal population of *B. thailandensis*, whereas other cells remained in a dividing or non-dividing state during exposure to phage. These new data are in line with recent observations by Attrill *et al.*, who reported the emergence of different phenotypes of *E. coli* during phage T4 predation, accompanied by a halving of the population through lysis with significant isolated survival (47). However, we did not observe significant isolated survival of *B. thailandensis*, suggesting that *B. thailandensis* and *E. coli* might adopt different phenotypic strategies to survive phage predation. We should also acknowledge that as a temperature-dependent lysogenic phage (27) infection with the ampunavirus ΦBp-AMP1 may be affected by lysogeny, and existant temperate phages may affect the observed interactions. We conducted our experimentation at temperatures in which ΦBp-AMP1 is known to be strictly lytic (27,28) and did not observe spontaneous cell lysis in the presence of the lysogen-activating antibiotic ciprofloxacin (48), thus suggesting that lysogeny did not affect the results discussed herein.

The interactions between bacteria, antibiotics and phage are complex and difficult to predict and may be affected by the environmental structure and the antibiotic mode of action, with antibiotics that share the same cellular target interacting differently with the same phage (14,18,31,49). We found that these interactions are further complicated by cellular heterogeneity within a putatively clonal population of *B. thailandensis* and that their study necessitates the development and use of approaches that allow single-cell resolution. Using such an approach we discovered that in the presence of a sub-inhibitory concentration of trimethoprim and phage, a large subpopulation of *B. thailandensis* performs doubling albeit at a slow rate and the subpopulation that lyses is reduced compared to phage alone leading to an antagonistic effect of trimethoprim on phage. These data are in line with previous bulk analysis showing that trimethoprim inhibits the synthesis of dihydropteroyl hexaglutamate, a vital molecule in the baseplate assembly of T-even and *Pseudomonas* phages, resulting in non-virulent phage progeny (50,51). We also found that a sub-inhibitory concentration of ciprofloxacin increased the frequency of phage-induced lysis of *B. thailandensis*. Interestingly, we found cells that were continuously dividing during phage predation to display a reduced cell size. The generation time was unaffected in the small-cell phenotype and untreated bacteria divided with the same frequency as long-term phage-treated bacteria. Campey *et al.* recently demonstrated that *E. coli* cells double while shrinking to overcome high ciprofloxacin stress in synthetic refuges (24), suggesting a potential double-while-shrinking strategy of *B. thailandensis* to overcome phage stress.

## Conclusion

*B. pseudomallei* is notoriously resistant to antibiotics, can cause recurring and chronic infection and often fatal disease. Its ability to hide from the immune system, antibiotics or phage in spatial refuges, such as within biofilms, is a major healthcare concern. Here we show that treatment of *B. thailandensis* with antibiotics or phage in synthetic refuges can be further complicated by cellular heterogeneity with the emergence of different phenotypes. We further show that a sub-inhibitory concentration of ciprofloxacin in combination with phage decreases cell size and increases the subpopulation of lysing or non-dividing *B. thailandensis*, whereas a sub-inhibitory concentration of trimethoprim in combination with phage delays cell growth but reduces the frequency of cell lysis. These findings can help inform the formulation of rational phage-antibiotic combination therapies to treat antibiotic-resistant infections caused by *B. pseudomallei*.

## Data availability

The data generated during this study and used in the figures are included in Supplementary files S1-S7.

## Author contributions

Conceptualization, S.P.; methodology, S.K., M.L.F., U.L., S.K., S.H., K.T.A., M.A.T.B. and S.P.; formal analysis, S.K., M.L.F., U.L., S.K., S.H., K.T.A., M.A.T.B. and S.P.; generation of figures, S.K. and S.P; investigation, S.K., M.L.F., U.L., S.K., S.H., K.T.A., M.A.T.B. and S.P.; resources, K.T.A., M.A.T.B. and S.P.; data curation, S.K. and S.P.; writing – original draft, S.K. and S.P.; writing – review & editing, S.K., M.L.F., U.L., S.K., S.H., K.T.A., M.A.T.B. and S.P.; visualization, S.K. and S.P.; supervision, S.P., M.A.T.B. and K.T.A.; project administration, S.P.; funding acquisition, U.L., M.A.T.B., K.T.A. and S.P..

## Acknowledgments

This work was supported by the BBSRC and the EPSRC through two grants awarded to S.P. and U.L. (BB/V008021/1, EP/Y023528/1). This work was further supported via the JPIAMR project ERADIAMR (MR/Y033892/1) awarded to S.P.. S.K. was supported by a QUEX PhD studentship awarded to S.P., M.A.T.B. and K.T.A. KTA also gratefully acknowledges the financial support of the EPSRC (EP/T017856/1). The funders had no role in study design, data collection and analysis, decision to publish, or preparation of the manuscript. The views expressed are those of the authors. For the purpose of open access, the authors have applied a ‘Creative Commons Attribution (CC BY) licence to any Author Accepted Manuscript version arising from this submission.

